# Differential nuclear import sets the timing of protein access to the embryonic genome

**DOI:** 10.1101/2021.10.18.464816

**Authors:** Thao Nguyen, Eli Costa, Tim Deibert, Jose Reyes, Felix Keber, Michael Stadlmeier, Meera Gupta, Chirag K. Kumar, Amanda Amodeo, Jesse C. Gatlin, Martin Wühr

## Abstract

The development of a fertilized egg to an embryo requires the proper temporal control of gene expression^1-6^. During cell differentiation, timing is often controlled via cascades of transcription factors (TFs)^7,8^. However, in early development, transcription is often inactive, and many TF levels are constant, suggesting that unknown mechanisms govern the observed rapid and ordered onset of gene expression^9^. Here, we find that in early embryonic development, access of maternally deposited nuclear proteins to the genome is temporally ordered via importin affinities, thereby timing the expression of downstream targets. We quantify changes in the nuclear proteome during early development and find that nuclear proteins, such as TFs and RNA polymerases, enter nuclei sequentially. Moreover, we find that the timing of the access of nuclear proteins to the genome corresponds to the timing of downstream gene activation. We show that the affinity of proteins to importin is a major determinant in the timing of protein entry into embryonic nuclei. Thus, we propose a mechanism by which embryos encode the timing of gene expression in early development via biochemical affinities. This process could be critical for embryos to organize themselves before deploying the regulatory cascades that control cell identities.

**Graphical abstract:** 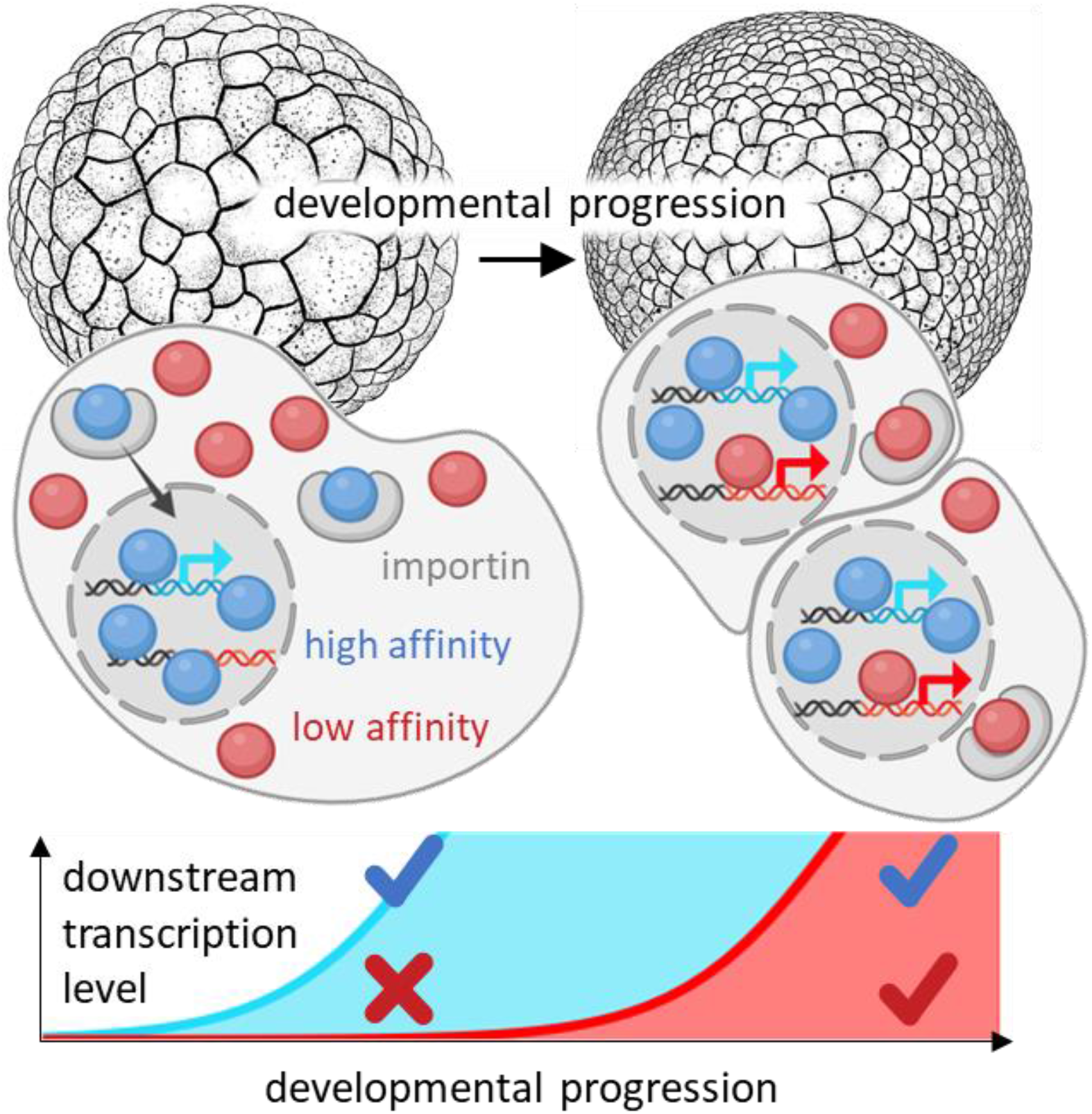

## Main Text

The oocyte is an exceptionally large cell (∼1.2 mm diameter in the frog *Xenopus laevis*) and in many species contains a proportionally large nucleus (Fig. 1a)^10-13^. After fertilization, the nucleus becomes tiny compared to the egg, and the nucleocytoplasmic volume ratio (NCV-ratio) drops by more than four orders of magnitude (Fig. 1a, b, Extended Data Fig. 1a)^14,15^ Most nuclear proteins are released into the cytoplasm during this drastic change in nuclear volume, but the total amount and composition of canonical nuclear proteins change little (Fig. 1c, Extended Data Fig. 1b, Supplementary Table 1)^9^. During the rapid early cleavage cycles that divide the single egg into thousands of cells, DNA and total nuclear volume increase approximately exponentially (Fig. 1b)^14^, and maternally deposited nuclear proteins from the oocyte nucleus are likely reimported into the newly forming nuclei. Whether all nuclear proteins from the oocyte re-enter the embryonic nuclei simultaneously or if there is a sequential order for nuclear import is unclear. What is known is that once the NCV-ratio reaches a critical value, embryos initiate several essential cellular activities, including the onset of zygotic transcription and cell movement^3,5,16-22^. This zygotic genome activation (ZGA) is an ordered process starting with transcripts dependent on Pol III followed by a specific temporal sequence of Pol II-dependent transcripts^19,23-28^. At later stages, the sequenced onset of transcription is often controlled via a sequenced expression of transcription factors (TFs)^7,8^. However, the inherent time delay between the transcription and translation of new genes is too large for it to explain the rapid and reproducible onset of different transcriptional events in the earliest developmental stages.

**Figure 1:**
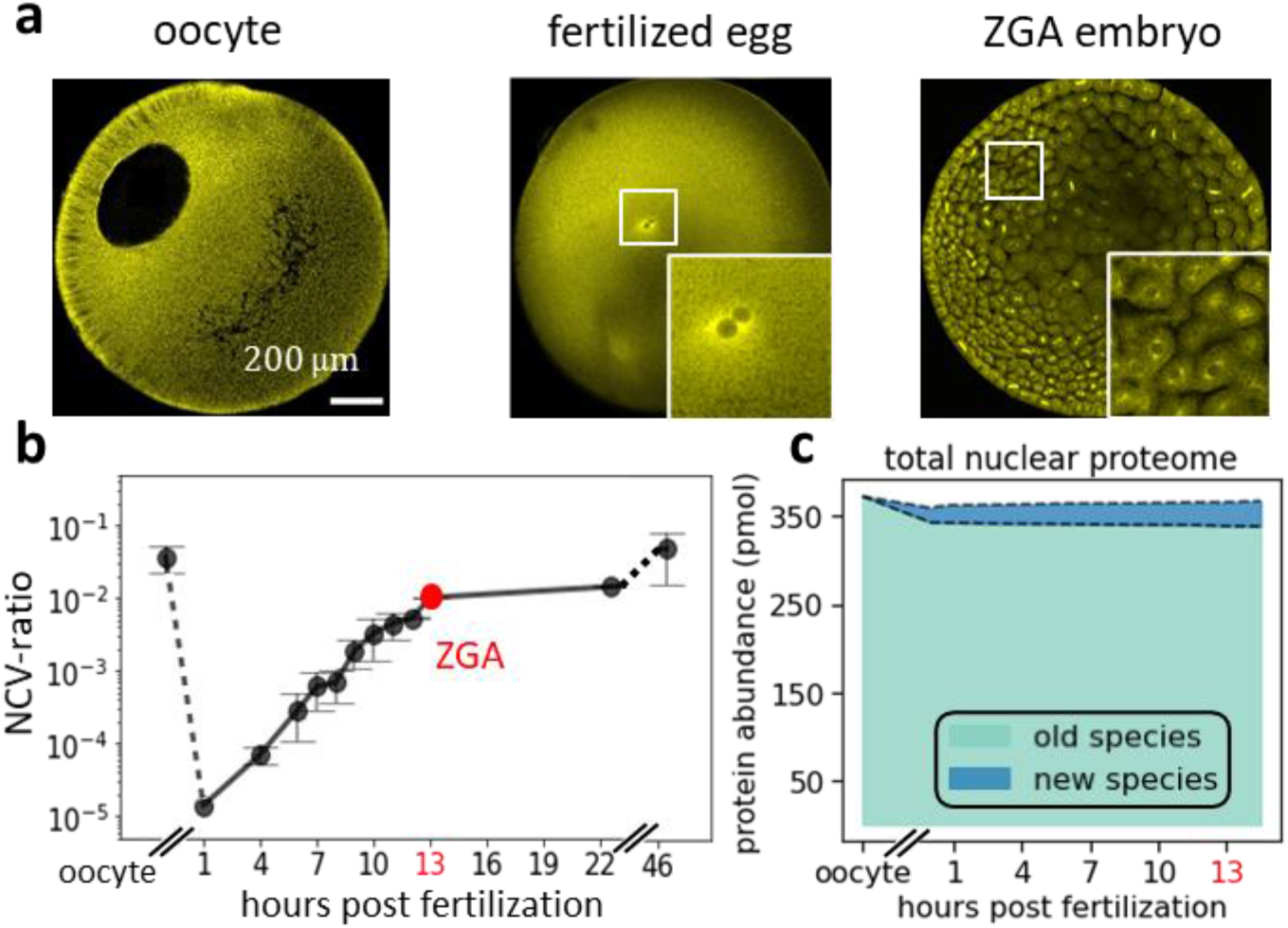
In early embryonic development, nuclear morphology changes drastically while nuclear proteome composition changes little. **a**, Immunofluorescence images (anti-tubulin) of early frog development show drastic changes in nuclear morphology. The large oocyte (∼1.2mm diameter) contains a proportionally large nucleus (∼400μm). After fertilization, the nucleus is only ∼30μm in diameter. Around ZGA, the embryo contains ∼4000 cells with ∼18μm diameter nuclei. **b**, The nucleocytoplasmic volume ratio (NCV-ratio) in early frog development, quantified based on micrographs. The NCV-ratio drops ∼10,000-fold from oocyte to fertilized embryo before increasing exponentially during early cleavage stages. After the ZGA, the NCV-ratio gradually increases approaching the value in the oocyte. Error bars indicate standard error. Embryos develop at 16°C. **c**, Quantification of the fraction of nuclear proteome replaced by new protein species from oocyte to ZGA. Despite the drastically increasing NCV-ratio from the fertilized egg to ZGA, approximately 3% of the nuclear proteome is replaced by newly synthesized canonical nuclear proteins (main contributors are high abundant histones, which increase ∼2-fold).

Furthermore, many TFs are often maternally deposited and show approximately constant expression levels (Extended Data Fig. 1c, Supplementary Table 1)^7,9^. Thus, it is likely that mechanisms beyond TF cascades are responsible for the sequential onset of gene expression in early development. We postulated that early embryos leverage their changing nuclear morphology by ordering the reimport of nuclear proteins such as TFs to time the onset of rapid downstream events.

### Proteins sequentially enter embryonic nuclei

To obtain insight into the times at which proteins gain access to the genome, we measured changes in the nuclear proteome over developmental progression in *Xenopus laevis* embryos. Using a novel protocol involving rapid filtration that allowed separating nuclei from other compartments, particularly mitochondria (Extended Data Fig. 2), we could enrich the nuclei at various embryonic stages and determine the nuclear fraction of each protein (FN) by quantifying its signal in the supernatant and the flow-through via multiplexed proteomics (Fig. 2a)^29,30^. We determined when half of the protein had entered the embryonic nuclei using a sigmoidal fit of the protein’s FN over time (T_embryo1/2_). Comparing T_embryo1/2_ across ∼2k canonical nuclear proteins, we found that the timing of nuclear import varied widely (Supplementary Table 2, Extended Data Fig. 3e). Extended Data Fig. 3a shows the FN data with sigmoidal fits for three example proteins (Nupl2, Smad2, Polr2b) with T_embryo1/2_ ranging from 12 hours to 31 hours, a representative spread for nuclear proteins (we define the time of fertilization as t=0).

**Figure 2:**
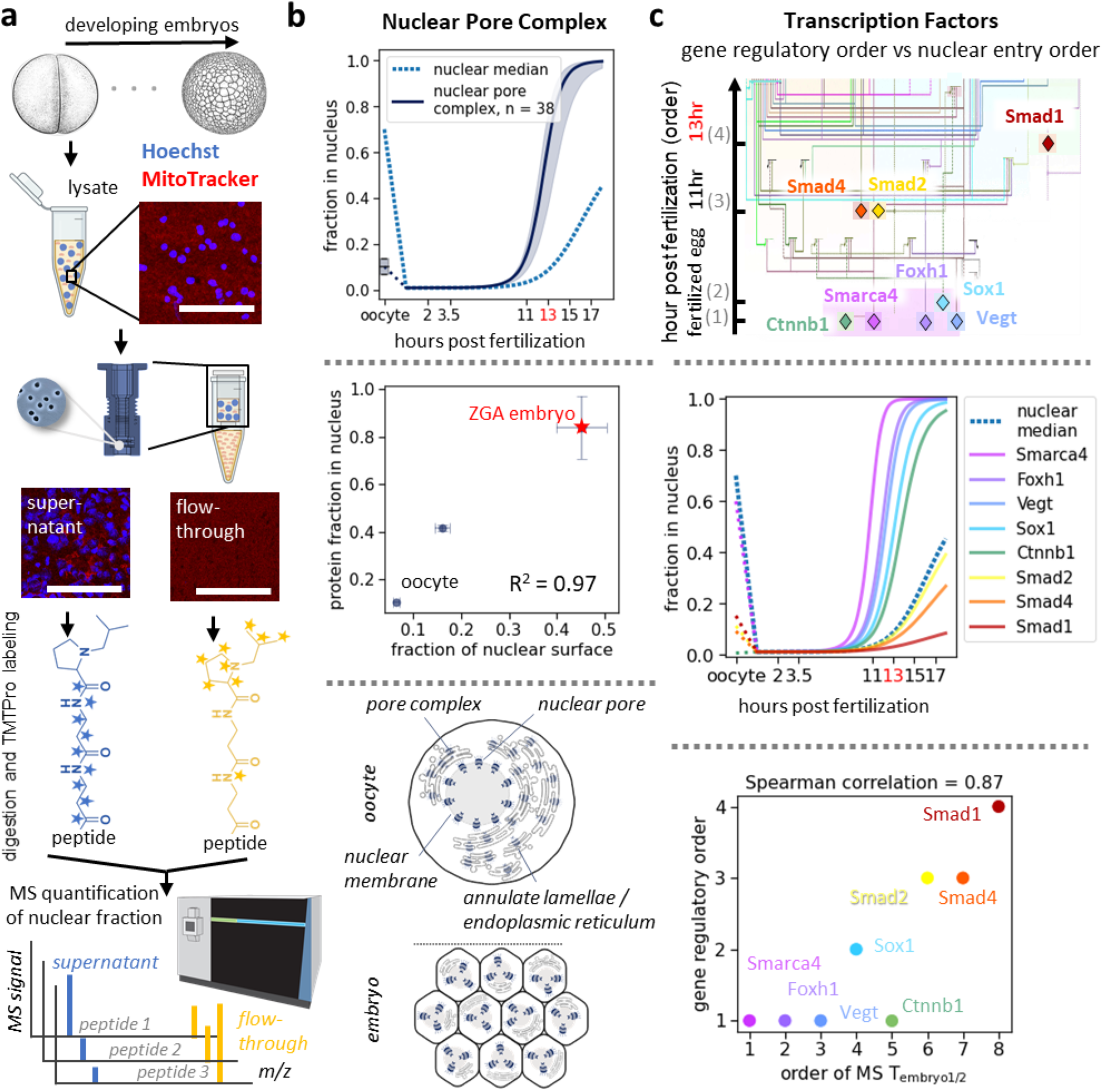
In early development, proteins enter nuclei sequentially correlating with the onset of their nuclear functions. **a**, Quantification of nucleocytoplasmic partitioning of the proteome in early development^54,55^. We collected embryonic lysate at various stages of development. To enrich the nuclei, we filtered the lysate through membranes retaining nuclei in the supernatant. Shown at each experimental step are Hoechst (blue) and MitoTracker (red) stained epifluorescence images of the lysate, supernatant, and flow-through. The supernatant and flow-through fractions are digested into tryptic peptides, labeled with isobaric tags, and subjected to accurate multiplexed proteomic analysis. **b**, Change in nucleocytoplasmic partitioning for nuclear pore complex (NPC) subunits corresponds to change in total nuclear surface area. Top: The blue dashed line shows the median nuclear titration of oocyte nuclear proteins (shaded area shows 50% spread of NPC subunits). Middle: Quantification of the nuclear fraction of NPC proteins via MS shows a good agreement with the change of nuclear surface quantified via immunofluorescence. We define the fraction of nuclear surface relative to embryos at 22.5 hours post fertilization. Bars indicate standard errors. Bottom: Our observations are consistent with previous electron microscopy studies indicating that NPC subunits are stored in endoplasmic reticulum membranes embedded with pore complexes in the oocyte, called annulate lamellae ^34,35^. **c**, Timing of nuclear import of TFs in the mesendoderm gene regulatory network corresponds to the timing of their activation. Top: Transcriptional activation order in early *Xenopus* embryos (Adapted drawing from Charney et al.^7^). Middle: MS-quantification of nuclear import in early development for these TFs. Bottom: Scatter plot of rank between the measured nuclear entry T_embryo1/2_ by MS and the reported temporal gene regulatory network shows strong agreement (Spearman correlation of 0.87, p-value = 0.005).

Validating the functional significance of our measurements, we found that the subunits of a given protein complex typically enter the nucleus at similar times. For example, although three different DNA repair complexes (the Fanconi anaemia complex (T_embryo1/2_ = 12 hours), the homologous recombination (T_embryo1/2_ = 20 hours), and the nonhomogeneous DNA end-joining repair complex (T_embryo1/2_ = 35 hours)) enter nuclei at different times (t-test p-value of 0.02), the subunits within a single repair complex arrive together (Extended Data Fig. 3b). Our observation of delayed sequential nuclear entry suggests that separating repair enzymes from DNA might contribute to the previously observed suppression of DNA repair during the rapid early cleavage cycles^31,32^. Nonetheless, we measured a few interesting cases where individual subunits within a complex enter the nuclei at different times. For example, we found that the essential core subunits of the origin of replication complex (Orc1-5) uniformly enter the embryonic nuclei early (T_embryo1/2_ = 12 hours), whereas a non-essential component (Orc6) enters at a much later time T_embryo1/2_ = 21 hours (Extended Data Fig. 3c)^33^.

Our protocol allows following proteins associated with the nuclear surface as well as proteins resident in the nucleus itself. We find that the nuclear pore complex proteins essential for entry of nuclear resident proteins retain a remarkably constant relationship to nuclear surface area. In oocytes, nuclear pore complex (NPC) proteins localize predominantly in the cytoplasm of oocytes (Fig. 2b, Extended Data Fig. 3d). In fertilized embryos the NPC proteins rapidly incorporate into the exponentially increasing nuclear surfaces. The nuclear accumulation of NPC subunits follows the corresponding changes of nuclear surface area in early development (R^2^ = 0.97). Specifically, we measured a 7.1-fold increase in the nuclear surface from the oocyte to embryo at the stage of zygotic gene activation (i.e., ZGA embryo), corresponding to an 8.1-fold increase in the nuclear accumulation of NPC proteins (Fig. 2b). These findings are consistent with previous electron microscopy studies showing that extra nucleoporins are stored in annulate lamellae in the oocyte cytoplasm before being incorporated in the embryo’s increasing nuclear surfaces (Fig. 2b bottom)^34-36^. These results suggest that import capacity per nuclear surface remains approximately constant during the cleavage divisions, despite the changing pattern of nuclear protein imported.

### Ordered entry of nuclear proteins may explain the timing of downstream activity

Our proteome-wide investigation of the changes in the nuclear proteome during early development reveals that the composition of the embryonic nucleus is dynamic. The ordered access of proteins to the genome could affect the timing of their nuclear functions. We next explored how differential nuclear import might relate to the control of RNA transcription. In frogs and many other species, transcription is inhibited during the early cleavages^3,19,20^ and is initiated only when the exponentially increasing DNA has titrated out a transcription inhibiting factor from the cytoplasm. DNA replication factors and histones have been proposed to be the titrated molecules in this model^21,22^. Indeed, we found these proteins were among the earliest proteins to enter the nuclei (Extended Data Fig. 3f). These factors become depleted from the cytoplasm around the ZGA, such that their concentrations begin to decline in the exponentially increasing nuclei. It has been proposed that the lower concentration of DNA replication factors slows down DNA replication, thereby enabling the onset of the very first transcripts^22^. Lowering nuclear histone concentrations are believed to change chromatin state, allowing transcription to start^21,37-41^.

Even though the embryo can transcribe at the ZGA, only Pol III transcripts are initially observable, followed by a clearly ordered sequence of Pol II transcripts^7,19,26,27^. We investigated whether sequential nuclear import could explain this ordering. We found that Pol III subunits enter embryonic nuclei comparatively early (T_embryo1/2_= 15 hours), while Pol II subunits are delayed (T_embryo1/2_ = 30 hours) (Extended Data Fig. 3g). This timing agrees with the well-established observation that Pol III transcripts (tRNA) are the very first to be transcribed in early development, followed by Pol II transcripts (snRNA and mRNA) (Extended Data Fig. 3g)^19^. Among Pol III transcripts, we observed a further correlation between the entry of TFs and the transcription of their downstream targets (Extended Data Fig. 3h). Gtf3c1-5 are direct transcription factors of tRNA, while Gtf3a is of 5S rRNA and 7S rRNA. Gtf3c1-5 enter the nucleus earlier than Gtf3a, which results in the appearance of tRNA before 5S rRNA and 7S rRNA^19^.

Subsequently, we investigated how the nuclear entry of specific maternally deposited Pol II TFs related to their downstream functions. Charney et al. (2017) described a gene regulatory network for maternal TFs active from fertilization through early gastrulation (Fig. 2c)^7^. We found that these TFs do not change their total expression levels in early development (Extended Data Fig. 1c) yet enter embryonic nuclei at vastly different times. Indeed, the order of their nuclear entry is highly predictive of the timing for their downstream gene activities, with a Spearman correlation of 0.87 (p-value = 0.005) between our measurements of TFs entry into the nucleus and the order in which they activate transcription in embryogenesis (Fig. 2c bottom). Thus, our analysis can explain not only why Pol III transcripts precede Pol II transcripts, but why individual Pol II transcripts arise in sequence despite their expression levels remaining constant.

### Importin affinities correlate with the timing of nuclear import

Next, we investigated the molecular mechanisms responsible for the differential timing of protein nuclear entry. Proteins could accumulate in the nucleus via active transport through binding to importins, proteins responsible for nuclear import, or by sequestration into the nucleus, e.g., by binding to DNA^15,42^. Differential protein affinities to either DNA or to importins might result in ordered nuclear import. We found that DNA affinity was poorly predictive of nuclear entry time (Extended Data Fig. 4a, b), suggesting that active transport may play a more significant role among the two mechanisms. Though the early embryo is known to express at least nine different importins, we focused on the most abundant, importin β (Kpnb1), and its canonical adapter, importin α1 (Kpna2) ^43,44^. We aimed to quantify the proteome-wide binding affinities of proteins to importins by exposing frog egg lysate to varying amounts of active importin β, controlled by different amounts of added RanQ69L. This constitutively active mutant mimics the GTP-bound form (Fig. 3a)^45^. Ran-GTP leads to conformational changes in importin β that induces the release of its substrates^46^. Following the addition of importin α1 and one-hour incubation in frog egg lysate, we then isolated importin β via affinity purification. Using quantitative proteomics, we measured the abundance of co-isolated proteins and their sensitivity to increasing amounts of RanQ69L (Fig. 3a). We integrated the information of our triplicate measurements by projecting the values on a single dimension, which we call importin affinity proxy, determined by cross-validated canonical correlation analysis (Fig. 3b, Supplementary Table 3)^47^.

**Figure 3:**
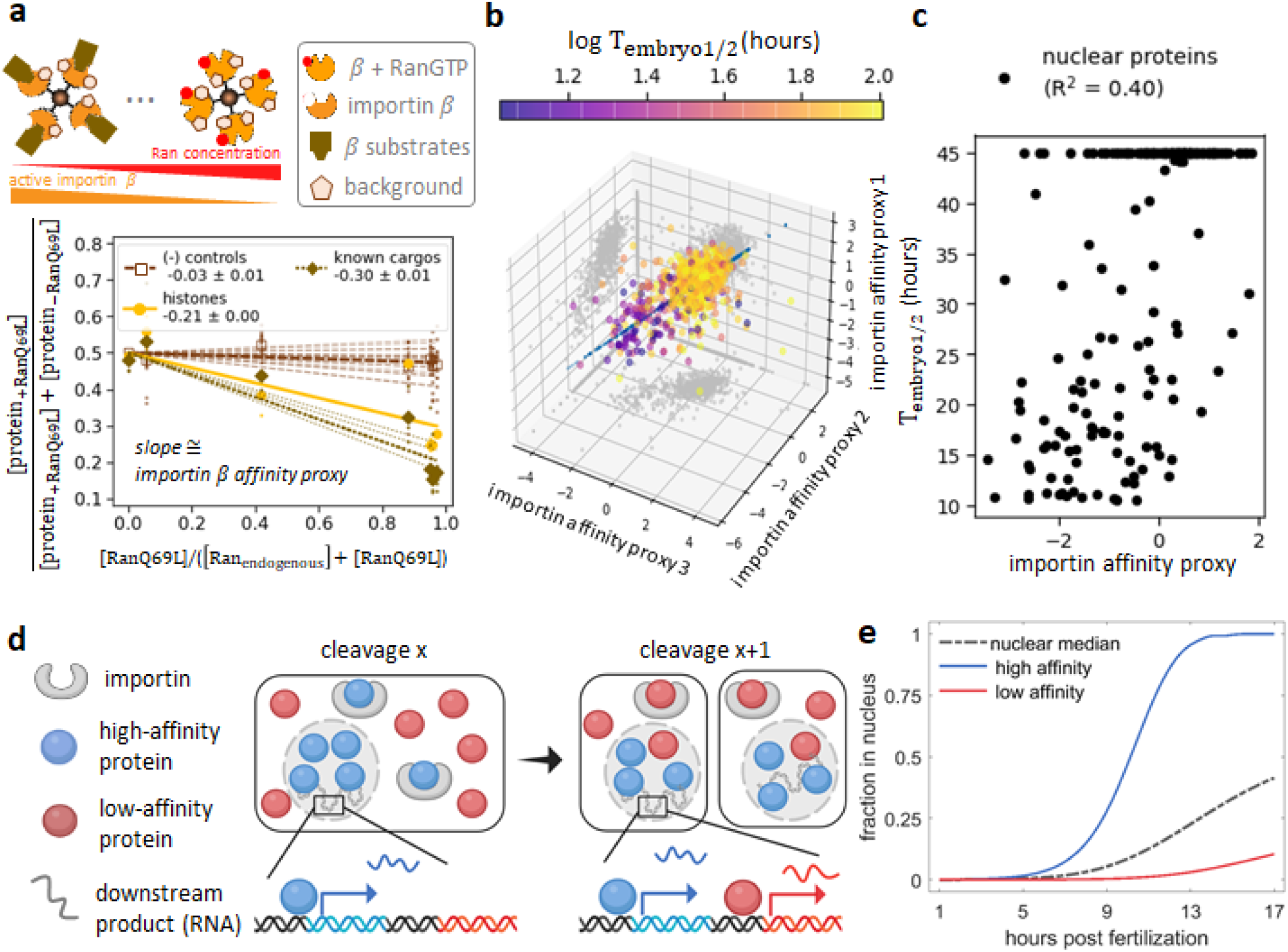
The affinity of proteins to importin contributes significantly to their ordering of nuclear entry in early development. **a**, Estimation of proteome-wide affinity to importin β. We measure the fraction of the MS signal coming from a condition with RanQ69L compared to the control. The MS signals of known importin β substrates^56^ including histones decrease with increasing RanQ69L concentration. Large dots represent the median protein fraction of a protein subgroup at each RanQ69L concentration, while small dots represent measurements for individual proteins. We applied a linear fit for each protein with a fixed y-intercept and use the slope as a proxy for a protein’s affinity to importin. **b**, Scatter plot of triplicate affinity proxy measurements from experiments outlined in **(a)**. We integrate these measurements on a dimension using cross-validated canonical correlation analysis ^47^. **c**, Importin affinity can explain a significant fraction of the timing of nuclear entry in early development. The scatter plot shows T_embryo1/2_ versus importin β affinity proxy. The observed Pearson correlation suggests that importin affinities can explain > 40% of the variance of timing of nuclear entry in early embryonic development. **d**, Schematic of our proposed model in which the differential affinity of proteins to importins controls the timing of genomic access in embryonic development. A high affinity protein titrates into the nucleus faster than a low affinity protein, resulting in the corresponding DNA access of proteins. For proteins associated with transcription, this determines when certain transcriptional products appear. **e**, Simulation of the model proposed in (**d**). We model competitive binding of substrates with varying affinity to a limited number of importin. The proposed model provides a simple explanation for the timing of protein access to the embryonic genome in early development.

When we plot the T_embryo1/2_ against the importin affinity proxy, we observe an R^2^ of 0.40 (p-value = 5e- 27) for canonical nuclear proteins (Fig. 3c). The correlation improved (R^2^ = 0.48) on including our DNA affinity measurements (Extended Data Fig. 4c). These R^2^ values are likely a severe underestimation of the actual contribution as they do not consider experimental noise in the embryo proteomics or the pulldown experiments^48^. Nevertheless, our ability to explain at least 48% of the observed variance for the timing of nuclear entry from these two simple assays is remarkable, especially considering that nuclear import in early embryos is undoubtedly more complicated than implied with these assays.

### A simple model explains the timing access on the genome in early embryos

Collectively, our data indicate that nuclear proteins enter nuclei at different stages of early development and that the timing of this entry correlates with affinity to importin. Therefore, we postulate that the limited amount of importin in cells (at ∼1.5µM, compared to ∼320µM of all protein with a predicted nuclear localization signal) binds predominantly to the highest affinity substrates (Fig. 3d, Supplementary Table 4)^49,50^. These substrates are then preferentially imported into early nuclei. After high-affinity substrates are depleted from the cytoplasm and imported into the increasing nuclear volume, importins become available to lower affinity substrates, and those proteins can then enter the nuclei. To formalize this hypothesis, we developed a simple model for competitive binding of substrates with varying K_D_, to limiting amounts of importin. Furthermore, we assume that nuclear import flux is partitioned among substrates based on their relative binding to importin. We derive the net nuclear import flux from immunofluorescence images (Extended Data Fig. 5a).

This straightforward model recapitulates the observed differential nuclear import of nuclear proteins well. It provides a quantitative framework for how the embryo could use substrate affinities to importin to determine the timing of protein’s import into the nucleus and, subsequently, their downstream nuclear functions (Fig. 3e). Interestingly, our model predicts that the concentration of nuclear proteins with high affinity to importin is high in the early nuclei and decreases as additional proteins are imported and the nuclear volume increases. Intermediate affinity substrates reach a maximal nuclear concentration at intermediate times before decreasing. Low affinity substrates are predicted to reach maximal nuclear concentration at late times (Extended Data Fig 5b). Finally, based on this model, we expect that the sequential nuclear protein import observed during developmental progression should also occur similarly in individual cell cycles after the nuclear envelope reforms.

### Nuclear entry in a single cell cycle following mitosis mimics the sequence in a developing embryo

During mitosis, the nuclear membrane breaks down and most nuclear proteins re-enter the cytoplasm. If importin affinities govern the global sequence of nuclear entry during the cleavage period, they should also govern re-entry on the shorter time scales following each round of mitosis. To test this possibility, we imaged the nuclear import following mitosis of nine GFP-labeled TFs observed in our embryo proteomics measurements in cell-free droplets prepared from egg lysates. The droplets were generated using a “T-junction” microfluidic device and contained demembranated sperm DNA, spontaneously forming nuclei (Fig. 4a left, Supplementary Video 1). Using time-lapse confocal microscopy, we then monitored the nuclear import of GFP-labeled proteins of interest (Fig. 4a right). To facilitate interexperimental comparisons of GFP-protein import rates, we used mCherry tagged with a nuclear localization signal (mCherry-NLS) as an import standard. From the relative nuclear-to-cytoplasmic intensity measurements, we extracted the half-time (T_droplet1/2_) of nuclear import for each protein and calculated the difference (ΔT_droplet1/2_) relative to mCherry-NLS observed in the same experiment (Fig. 4b). We observed strong agreement (Spearman correlation of 0.82, p- value = 0.007) between the order of nuclear protein import in embryos and the order of nuclear protein entry in cell-free droplets via imaging (Fig. 4c). This agreement holds despite the drastic changes in morphology and the potential differences in protein expression levels and post-translational modifications between the egg and the post-ZGA embryo. Our results suggest that differential nuclear entry could be used as a biological timing mechanism within each cell cycle. Additionally, we find that the nuclear protein concentration of early imported substrates (like Yy1) is initially high and decreases over time (Fig 4d). Protein Gtf2e2 reaches the maximum concentration at an intermediate stage. Lastly, late protein Gtf2b shows the highest concentration at the end of our measurements. These observations agree well with the predictions of our simple model for proteins with high, intermediate, and low affinity for importin (Fig 4e).

**Figure 4:**
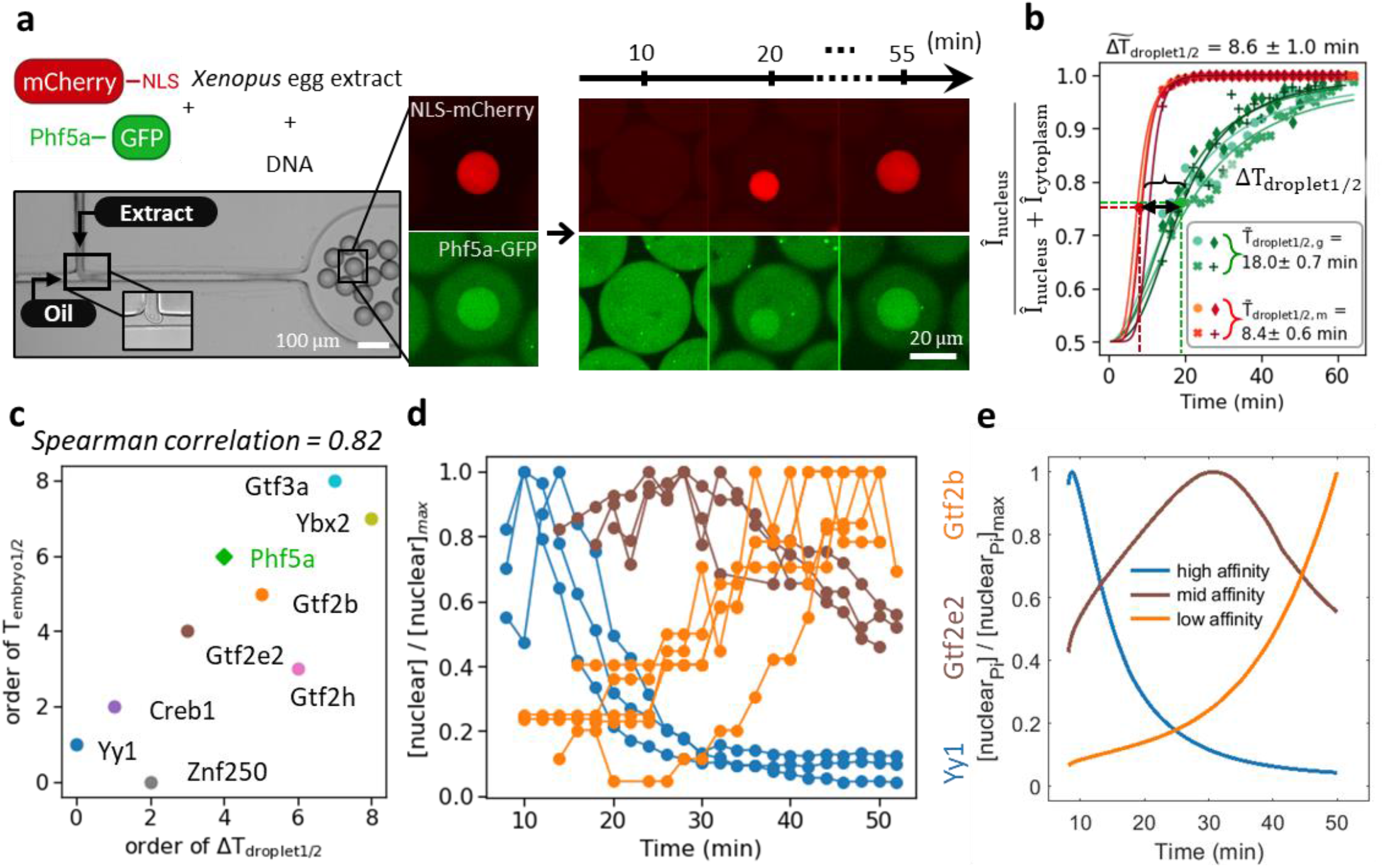
The temporal order of nuclear import in cell-free droplets observed via imaging recapitulates the nuclear entry order observed in the embryo proteomic assay. **a**, Left: Imaging of nuclear import in cell droplets. *Xenopus* egg extract doped with sperm DNA, which initiates formation of nuclei, and GFP-tagged protein of interest (here Phf5a) and mCherry-NLS were encapsulated in oil droplets with a microfluidic device. Right: We monitored nuclear import kinetics via fluorescence microscopy. **b**, We quantified the relative fluorescent signal intensity in nucleus and cytoplasm and fit the data with a sigmoid to extract the time (T_droplet1/2_) at which the relative intensity reaches half of its max value. To overcome extract variability, we calculate the import time difference (ΔT_droplet1/2_) between mCherry-NLS and the protein of interest. Markers represent the raw measurements. The different symbols represent different droplets; lines are sigmoid fits of corresponding droplets. From these experiments, we extract the median ΔT_droplet1/2_. **c**, Scatter plot of the order in nuclear import time (ΔT_droplet1/2_) from the cell-free assay and the order in T_embryo1/2_ for the nine TFs show strong agreement (Spearman correlation of 0.82, p-value = 0.007). **d**, Imaging results show the concentration of early titrator nuclear protein (Yy1) is high at the early stage and decreases over time, followed by Gtf2e2 and Gtf2b. **e**, In our import model, we predict that nuclear concentration of high affinity proteins (blue) is high in the early nuclei and decreases with the continuous import of additional nuclear proteins and the increasing nuclear volume. Nuclear proteins with lower affinities (brown then orange) will reach their highest nuclear concentration at some later times and in a sequence corresponding to their interaction strengths to importin. The imaging results are consistent with the model.[nuclear P_i_] is the nuclear concentration of protein *i* (*i* = blue, brown, orange) that is being evaluated.

## Discussion

Converting a fertilized egg to an embryo with a canonical body plan and hundreds of different cell types requires remarkable organization in space and time. While we have learned much about the spatial organization in early embryos, our understanding of embryo organization in time is substantially less refined^7,8,51^. Cascades of TFs provide natural order to the timing of newly expressed genes during cell differentiation ^8,52,53^. However, early development occurs at an astonishingly fast speed and is mostly transcriptionally silent. The inherent time delay between the transcription and translation of new genes is not consistent with the rapid and reproducible onset of different transcriptional events. Despite the drastic changes of embryonic morphology in early development, the proteome is remarkably constant (Fig. 1c, Extended Data Fig. 1b)^9^. For example, the TFs responsible for mesendoderm formation show approximately constant expression levels (Extended Data Fig. 1c)^7^.

Here, we find evidence that rapid early events in development might be ordered via biochemical affinities of maternally deposited nuclear proteins to importin. We find that proteins believed to inhibit the onset of transcription in early embryonic development partition to the nucleus early (Extended Data Fig. 3f). Subsequently, they are diluted with increasing nuclear volume. Once the embryo is permissive for transcription, the timing of protein entry into embryonic nuclei correlates strongly with the activation of their nuclear functions (Fig. 2c). We put forward a model, in which nuclear import is partitioned based on relative affinity to importin. This model predicts similar ordering for nuclear import during a single cell cycle. To test this model, we performed nuclear import assays in droplet encapsulated cytoplasm (Fig. 4). Furthermore, our model predicts that high affinity proteins decrease their nuclear concentration over time while medium and low affinity proteins reach maximal nuclear concentrations at later stages (Fig. 4e, Extended Data Fig. 5b). Our model can explain the loss of inhibitory effects on transcription before the ZGA for high importin affinity proteins like histones due to their lowering of nuclear concentrations with increased nuclear volume^21,22^. At the same time, the model can explain how lower affinity transcription factors enter the nuclei only at a later time, controlling the temporal onset of their downstream nuclear functions.

In this study, we quantify the temporal entry of ∼2k nuclear proteins. While we could only investigate a limited set of proteins for the timing of their downstream nuclear functions, it is very likely that the embryo widely uses the observed inherent timing mechanism. In addition, over 40% of the observed time-variance in nuclear import across all nuclear proteins was explained via differential affinities of the proteins for importin, despite quantifying affinities to only importin β using a crude and noisy biochemical assay. This predictive power suggests that this fundamental biochemical mechanism plays a crucial role in the temporal organization of developing early embryos that could set the stage before gene regulatory networks orchestrate cellular differentiation.

## Acknowledgments

We thank Matt Sonnett, Eyan Yeung, Lillia Ryazanov, Nick Treen for help and training and Eric Wieschaus, Mike Levine, Elizabeth Van Itallie, and members of the Wühr Laboratory for their comments and edits on the manuscript. We thank David Hill for access to the Xenopus ORFeome and Dirk Görlich, Thomas Güttler, and Sabine Petry for gifts of RanQ69L and importin plasmids. We thank James Pelletier for help with designing the filter holders. This work was supported by NIH grant R35GM128813 (MW), P20-GM113132 (AA), R01GM135568 (TD, JG), American Heart Association predoctoral fellowship 20PRE35220061 (TN), EMBO ALTF 601-2018 (FK), Princeton Catalysis Initiative (MW), Eric and Wendy Schmidt Transformative Technology Fund (MW), Harold W. Dodds Fellowship (MG), Princeton University’s Summer Undergraduate Research Program (EC, JR, CK), NSF MODULUS award 2052640 (TD, JG).

## Autor contribution

MW and TN conceptualized the study. TN and EC performed importin affinity experiments. TN and JR performed DNA affinity experiments. TD performed cell-free droplet experiments with JG’s supervision. TN and FK analyzed imaging data. TN and MS generated and analyzed mass spectrometry data. TN and MG developed the statistical framework. CK analyzed absolute abundance data. MW, AA, and JG provided funding. MW, JG, and AA supervised the study. TN, MW, and EC wrote the manuscript, and all authors helped edit the manuscript.

