## Supplementary material for "Differential nuclear import sets the timing of protein access to the embryonic genome"

### Extended Data Figures

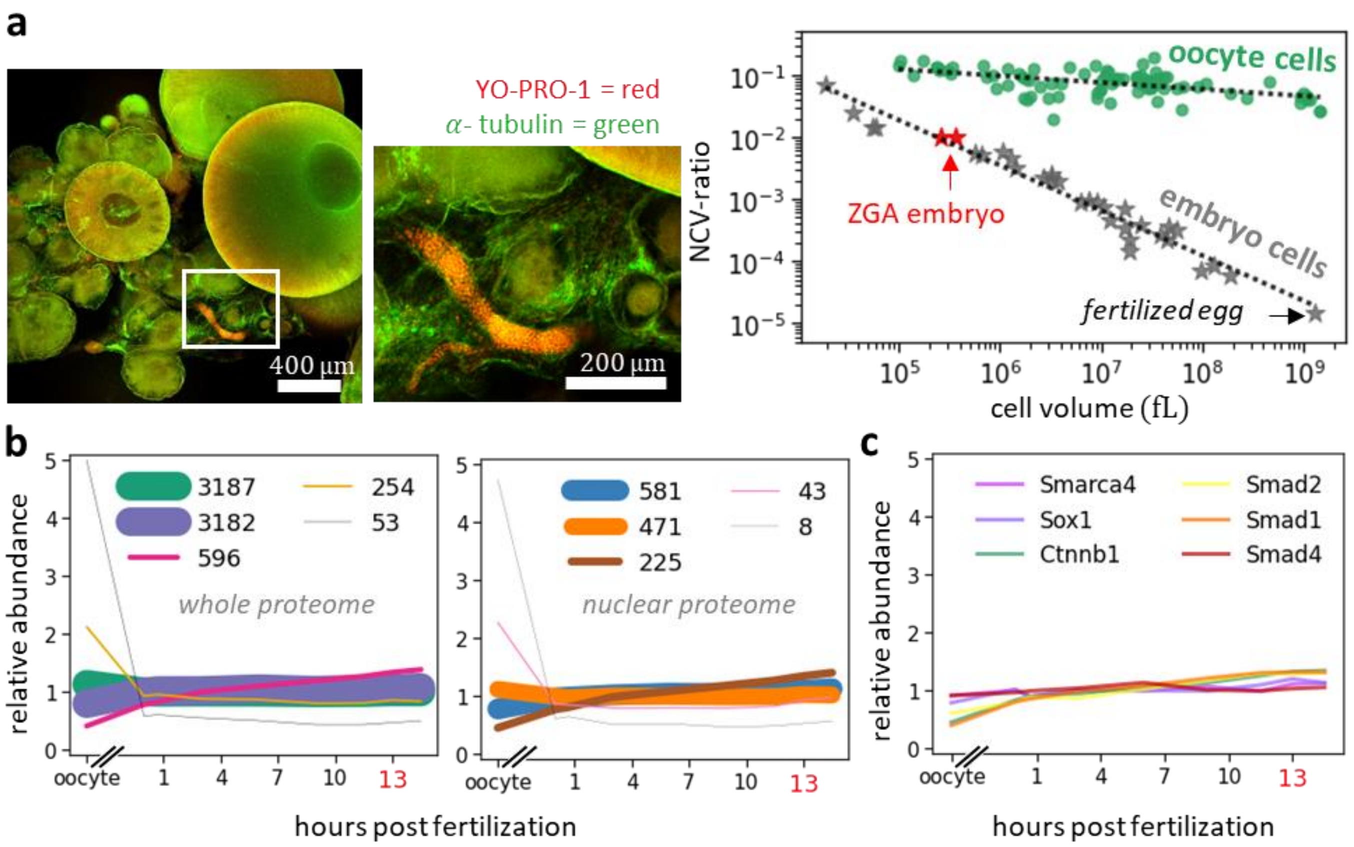


**Extended Data Figure 1: Changes in nucleocytoplasmic volume (NCV) ratio for cells in the frog ovary and early development.**

**a**, Left: Immunofluorescence (α-tubulin and Yo-Pro-1) of a frog ovary shows cells varying by 3-orders of magnitude in diameter (from ~10µm to ~1200µm). A zoomed-in section of the white framed box is shown to the right.

Right: Quantification of the NCV-ratio versus cell volume for maturing oocytes and early developing embryos. Compared to the rapidly changing embryos, the oocyte NCV-ratios remain approximately constant (decreasing by 2.3-fold as the cell volume increases ~500-fold (from ~2nL to ~1µL)) throughout oogenesis. The slight decrease might come from some cytoplasmic volume being excluded by the forming yolk platelets[1],[2]. In contrast, the NCV-ratios increases by ~34,000-fold in developing embryos from a 1µL fertilized egg to ~30pL cells at 46 hours post-fertilization (16°C).

**b**, k-means clustering (k=5) of changes in relative protein abundance of the entire proteome (left) and nuclear proteins (right) from the time series of the oocyte to ZGA embryo measured by multiplexed proteomics. The line thickness scales with the number of proteins within a cluster. While the NCV-ratio drastically increases from the fertilized egg to the ZGA, protein levels remain largely unchanged.

**c**, The detected transcription factors reported being involved in the gene regulatory network of the mesendoderm formation do not change their expression levels between fertilization to the ZGA[3].

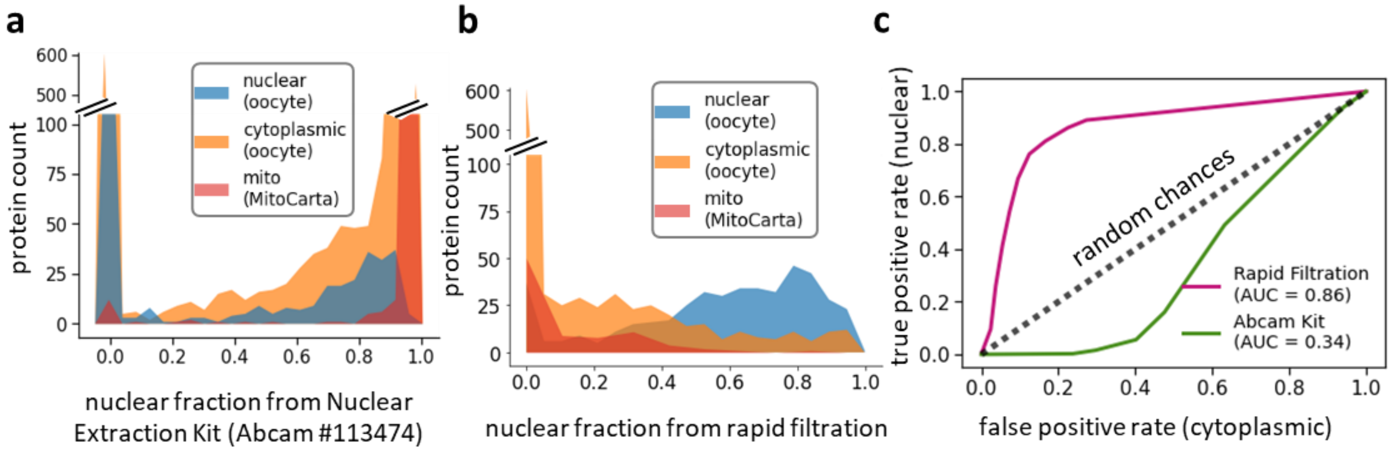


**Extended Data Figure 2: Rapid nuclear filtration outperforms nuclear isolation method based on differential sedimentation in quantifying the nucleocytoplasmic (NC) partitioning in early frog embryos.**

**a**, Nuclear extraction based on differential sedimentation poorly separates nuclear proteins from the proteins of other organelles. Shown is the histogram of nuclear fraction, quantified with multiplexed proteomics, after the fractionation of ZGA embryos using a commercial nuclear extraction kit (Abcam).

**b**, Our newly developed rapid filtration method is used to isolate nuclei and the nuclear fraction is quantified as in (**a**).

**c**, Receiver operating characteristics (ROC) for the measured nuclear fractions, comparing filtration and sedimentation-based nuclear enrichment methods. We use proteins that were measured nuclear in the oocyte as true positives and proteins that were measured cytoplasmic in the oocytes as false positives[4]. Measuring NC partitioning via rapid filtration (pink) appears superior to sedimentation (green) in early frog embryos.

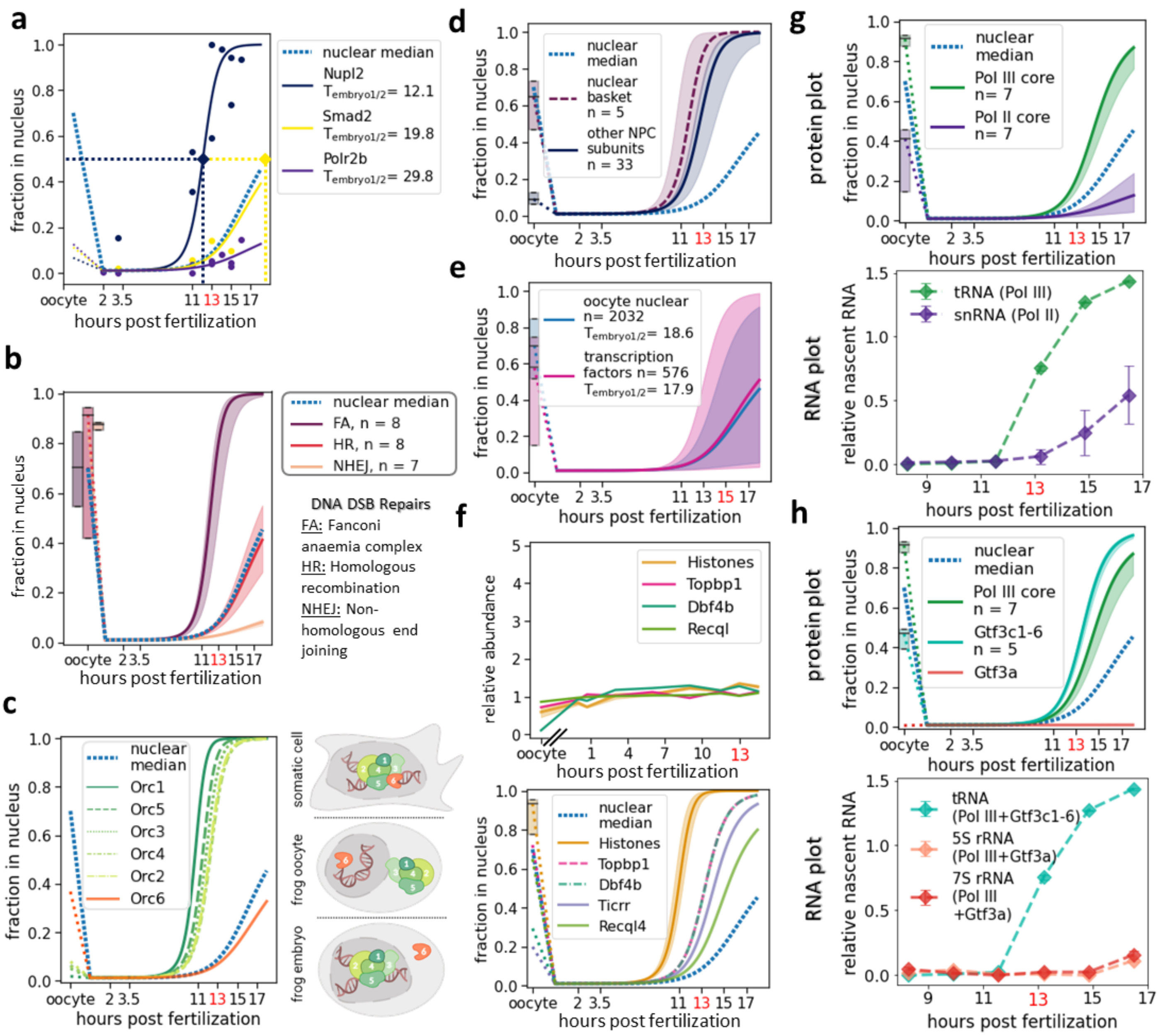


**Extended Data Figure 3: Quantitation of protein NC partitioning reveals sequential nuclear entry in early embryos.**

**a-h**, x-axis represents hours post-fertilization for an embryo that develops in 16°C. For complexes with multiple subunits, we represent the median nuclear titration pattern by a solid line and a 50% spread by shaded area with the corresponding color.

**a**, Example nuclear quantification for three proteins. The measured fractions of each protein in the nuclei at each stage are shown as dots. A sigmoid function (shown as solid lines) helps extract the time to the half max ($T_{embryo1/2}$). The three nuclear proteins exhibit drastically different nuclear entry times. For comparison, the median of nuclear proteins is shown (blue dash line).

**b**, Different DNA repair complexes enter embryonic nuclei at various times, yet their respective subunits enter simultaneously. Shown are nuclear entry quantifications for homologous recombination (HR), the nonhomologous DNA end-joining repair (NHEJ), and the Fanconi anaemia (FA). An apparent time delay of nuclear entry for some of the DNA repair complexes explains previous observations that early embryos under-correct or even bypass DNA repair to accommodate the fast-dividing time[5-7]. The oocyte nuclear fractions are shown in box plots with a median value and 50% spread.

**c**, Orc6 enters embryonic nuclei much later than other components of the origin of replication complex (ORC). While Orc1-6 localizes in somatic cells' nuclei, in the oocyte Orc1-5 are cytoplasmic and Orc6 is nuclear[4,8]. However, in early embryos, Orc1-5 rapidly enter the nuclei, presumably to accommodate for the core function in DNA replication processes in *Xenopus*[9]. The non-essential component Orc6 enters later. This is consistent with a previous observation of Orc6's cytoplasmic function in *Drosophila* and mammalian cells[10].
Right: A cartoon model summarizing the NC partitioning pattern of the ORC core subunits in the oocyte, embryo, and somatic cells.
**d**, The nuclear basket proteins show different NC partitioning in early development than other nuclear pore complex (NPC) proteins. While most NPC proteins are shared between nucleus and cytoplasm in the oocyte, the nuclear basket is mostly in the nucleus. Additionally, the nuclear basket also titrates into the nucleus slightly faster than the other subunits. Previous observations show that nuclear baskets are not part of the annulate lamellae, but tightly interact with the higher-order actin filamentous network that connects the NPC to chromatin in the oocyte nucleus[11].
**e**, Quantification of NC-partitioning as a function of developmental progression for ~2k nuclear proteins and ~600 transcription factors (TFs) quantified in our proteomics experiment [12-14]. At the ZGA, a significant fraction of nuclear proteins and TFs are still in the cytoplasm with a widespread of the $T_{embryo1/2}$.
**f**, Top: Protein abundance dynamics for key factors previously reported to be associated with the ZGA during development[15,16]. The protein abundance of these factors is largely constant from fertilization to the ZGA.
Bottom: These ZGA regulators are among the earliest proteins to titrate into embryonic nuclei. Shown are fits of individual ZGA regulators and the median fit of the core histones (H2a, H2b, H3, and H4) with a 50% spread of the four core histones and their isoforms.
**g-h**, The bottom plots of these panels are quantified from Newport and Kirschner's 1982 RNA gel of newly synthesized transcripts[17].
**g**, RNA polymerase III (Pol III) and II (Pol II) enter nuclei at different times of development, which correspond to the respective appearances of their first downstream transcripts.
Top: Proteomics data show that Pol III-specific subunits (green) titrate into nuclei before Pol II-specific subunits (purple).
Bottom: The first tRNA transcripts, transcribed by Pol III, and snRNA, transcribed by Pol II, show an identical order of appearance as the nuclear entry order of the polymerases[17].
**h**, Transcription factors titrate into the embryonic nucleus at different times which, for some, link to their downstream transcripts.
Top: Although the core Pol III titrates in the nucleus early, the nuclear entry times of its associated transcription factors vary. In our proteomics analysis, Gtf3c1-5 titrate in the nucleus before Gtf3a. Under Pol III, the transcription factors Gtf3c1-5 (and Pol III) are upstream of tRNAs, while Gtf3a (and Pol III) are upstream of 5S rRNA and 7S rRNA.
Bottom: Quantification of the newly transcribed RNA indicates that tRNA is transcribed much earlier than 5S rRNA and 7S rRNA, which corresponds to the nuclear entry order of their upstream transcription factors.

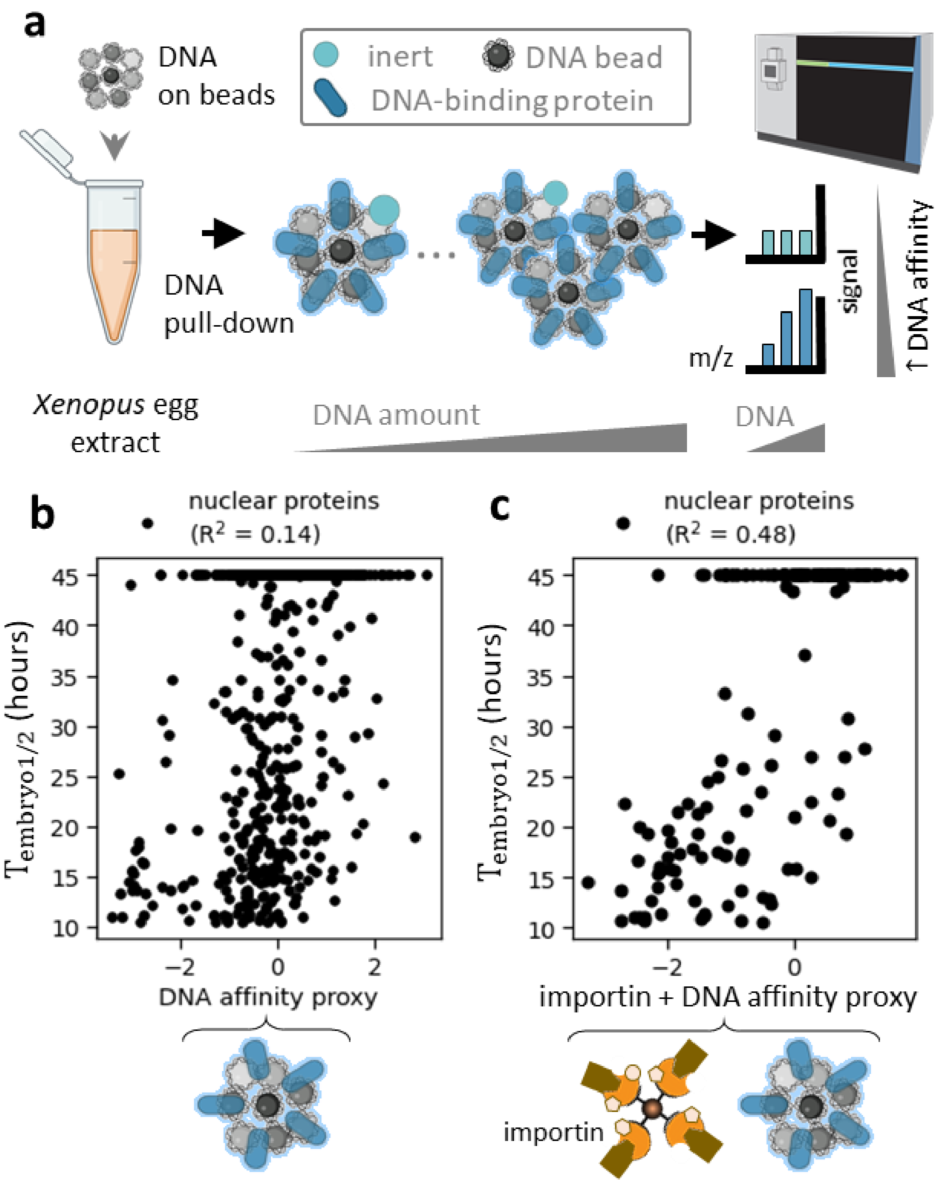


**Extended Data Figure 4: Quantification of proteome-wide affinities to DNA and their correlation with nuclear entry times in early embryos.**

**a**, Quantification of proteome-wide protein affinity to DNA. We exposed *Xenopus* egg lysate to magnetic beads that are covered with DNA. The pull-down is collected and subjected to MS quantification of the relative protein signal. The experiment was repeated with different DNA to lysate ratios. The DNA affinities at different ratios were projected on one dimension of the canonical coordinate space with cross-validation to produce a DNA affinity proxy for each identified protein[18].

**b**, Scatter plot of the projected DNA affinity proxy versus $T_{embryo1/2}$. The correlation suggests >14% of the nuclear import variance in the embryos can be explained by DNA affinity.

**c**, The combined DNA affinities and importin affinities projected onto a single dimension result in an improved correlation with nuclear entry time, suggesting that together DNA and importin affinities can explain >48% of the variance of observed timing of nuclear entry in early embryos. Shown is the scatter plot between $T_{embryo1/2}$ and the projected affinity proxy.

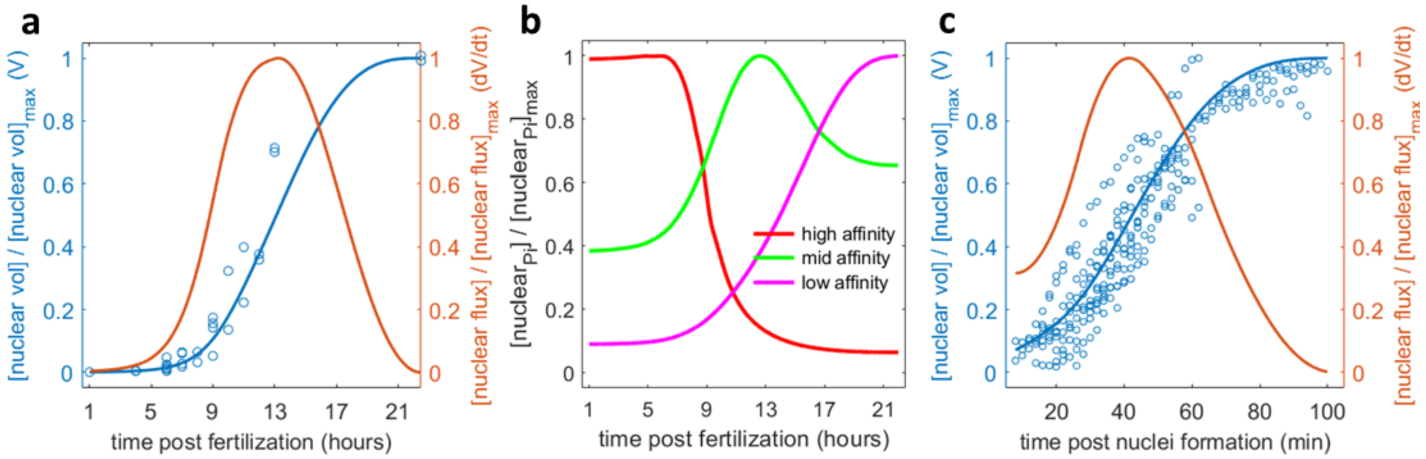


**Extended Data Figure 5. Changes of total nuclear volume in embryos and droplet assay and predictions of nuclear concentration for proteins with varying importin affinity in embryo.**

**a**, Total nuclear volume and nuclear protein flux as functions of time in developing *X. laevis* embryos. Each dot indicates measurements of total nuclear volume based on immunofluorescence from one embryo. A spline fit (blue curve) provides the functional form of the changes in nuclear volume over time. The time derivative of nuclear volume expansion is the nuclear flux due to nuclear import (orange curve). The maximum volume during this time normalizes the nuclear volume (y-axis left), and the nuclear flux is normalized by the maximum flux (y-axis right).

**b**, Our model predicts the changes in nuclear concentration over increasing embryonic nuclear volume for three proteins with different affinities to importin. The three proteins sequentially reach their maximal nuclear concentration from the highest affinity protein to the lowest affinity protein. After the maximum is reached, further nuclear volume increase leads to a decline in nuclear concentration.

**c**, Nuclear volume and nuclear flux as functions of time in oil encapsulated cytoplasmic droplets. Similar to (**a**), the nuclear volume per cell-free droplet is shown increasing with time in the raw imaging data (blue dots) and is fit by a spline function (blue curve). The function's time derivative is the nuclear flux over time (orange curve). T = 0 is the time when the cytoplasm is taken off ice and nuclear formation is initiated.

**Supplementary Information Guide**

**Supplementary table 1.** Results of quantitative proteomic measurement of the relative protein abundance over a developmental time series from the mature oocyte to pass the ZGA.

**Supplementary table 2.** Proteomics quantification of the half-times that proteins enter embryonic nuclei ($T_{embryo1/2}$) and nuclear fraction (NF) over a developmental time series.

**Supplementary table 3.** Proteomic estimation of importin β affinity, DNA affinity, and importin β + DNA affinity.

**Supplementary table 4.** Absolute abundance of proteins in the frog eggs reanalyzed with *X. laevis* protein Fasta file based on genome version 9.2[19].

**Supplementary movie 1.** The movie shows the formation of ~50µm diameter cell-free droplets of *Xenopus* egg extract in a continuous oil phase using a T-junction microfluidic device.

**Methods**

Oocyte, egg, and embryo collection

Mature *Xenopus laevis* females and males were purchased from Nasco and maintained by Laboratory Animal Resources at Princeton University. All animal procedures are approved under IACUC protocol 2070, reviewed in April 2021. For ovary collection, *Xenopus* females were euthanized in 0.1% aminobenzoic acid ethyl ester (Tricaine, MS222) (Sigma A-5040) and then sacrificed by pitching. For testes collection, an equivalent procedure was followed with male frogs. Females were ovulated with at least 6-month rest intervals. *X. laevis* oocytes, eggs, and testes were collected as previously described[20].

*Oocyte collection:* Female frogs were first primed by injecting 100 U of pregnant mare serum gonadotropin (PMSG) into the dorsal lymph sac at 3-60 days before the experiment to ovulate eggs. *Xenopus* females were sacrificed as described above on the day of the experiment. Ovaries were collected in a petri dish and cultured in an oocyte culture medium (1 L of OCM: 1 bag of Leibovitz's L-15 Medium powder (ThermoFisher Scientific #41300039), 8.3 mL Penicillin/Streptomycin, 0.67 g BSA) with the pH adjusted to 7.7 using NaOH and passed through a 0.22 µm filter[21]. We kept oocytes for up to a week and exchanged OCM daily. Alternatively, in later stages of the study, defolliculated oocytes from Ecocyte Bioscience US LLC were used.

*Egg collection:* At 16 hr. before egg collection, female frogs were injected with 500 U of human chorionic gonadotropin (HCG) and kept at 16°C in Marc's Modified Ringer's (MMR: 100 mM HEPES pH 7.8, 2 mM EDTA, 2 M NaCl, 40 mM KCl, 20 mM $MgCl_2$, and 40 mM $CaCl_2$)[22]. We collected eggs the next day in MMR buffer and sorted out pre-activated ones for further use.

*Embryo collection:* For *in vitro* fertilization, we first isolated testes from male frogs as previously described[20]. The testes were stored in OCM at 4°C and exchanged daily for up to one week of use. Then, we collected eggs from females onto Petri dishes by gently squeezing the frog. One quarter of one testis was used per 500 eggs by first crushing then mixing in the eggs using a sterile pestle. The mixture was incubated at room temperature (RT) for 5 minutes followed by another mix and an additional 5 minutes of incubation. Fertilization was then induced by flooding the eggs with 0.1× MMR. After about 30 min in 16°C, embryos uniformly rotated to face their pigmented halves upwards, indicating that fertilization succeeded. After this checkpoint, embryo jelly coats were removed by incubating with 2% L-cysteine in 0.1× MMR at pH 7.8 with NaOH for 2-5 min or until the jelly coats appeared to be removed. Embryos were then washed thoroughly with 0.1x MMR to remove any residual cysteine. Embryos develop in 0.1x MMR at 16°C until the desired time points.

We staged embryos based on Nieuwkoop and Faber[23].

Immunofluorescence of oocytes and embryos

The immunofluorescence procedure was performed essentially as described by Nguyen et al.[24]. Briefly, at stages of interest, ~20 embryos were arrested at interphase using cycloheximide, then collected and fixed with Methanol/EGTA for 24 hours. Embryos were then rehydrated using 25%, 50%, 75%, and 100% TBS (10 mM Tris–HCl, pH 7.4, 155 mM NaCl, and 0.65 g/L of $NaN_3$ to inhibit bacterial growth) and then in Methanol before bleaching with $H_2O_2$.

Embryos were re-submerged in TBSNB (TBS+ 0.1% Igepal CA-630, 1% BSA, 2% fetal calf serum). Samples were then incubated with α-tubulin (B-5-1-2) (Sigma T6074) that was pre-labeled with Alexa-488 using APEX™ Antibody Labeling Kits (Invitrogen) at a 1:200 dilution in the dark at 4°C for 12 hr., followed by washes in TBSNB for 24hr and two washes in TBS for 10 min. Finally, samples were dehydrated in Methanol and cleared by Murray's clear (2:1 Benzyl benzoate: Benzyl alcohol), before being mounted on a custom-made mounting slide for confocal imaging analysis[24].

Image analysis was performed on a laser scanning microscopy Zeiss 880 confocal microscope. The acquisition sequences were like previously published protocols[25]. For early-stage embryos, only a single focal plane that captured all cells was acquired. For later-stage embryos with multiple cells, a stacked image at 7.3µm step size was acquired along the animal and vegetal axis. Images were analyzed using ImageJ-win64 version 1.8.0.
To quantify the nucleocytoplasmic volume ratios, we assumed that embryos are rotationally symmetric and quantified nuclear volume for the embryos' representative sector. We assumed nuclei to be spherical and derived the volume for each nucleus from its measured diameter. We calculate average cell volume by dividing embryo volume by previously reported or newly measured cell numbers[23,26].

Nuclear filtration

To make embryo extract, embryos at the 2-cell, 4-cell, cleavage 10,11,12 (the ZGA), 1- hr. post-ZGA, and 3-hr. post-ZGA developmental stages were collected and prepared into lysate mostly as previously described[27]. Briefly, around 200 embryos were collected per time point. Embryos were first arrested in the interphase by incubating in 150ug/mL cycloheximide for 1hr. and then transferred into a standard 0.2mL PCR tube filled with ELB (250mM sucrose, 50mM KCl, 2.5 mM MgCl2, 10mM HEPES pH 7.8 with KOH) + 1ug/mL LPC + 1ug/mL cytochalasin D + 100ug/mL cycloheximide. Embryos were packed and crushed at the maximum accessible speed (11,600g) on a tabletop centrifuge. The cytoplasmic layer was withdrawn with a 23-gauge needle. The embryonic extract was supplemented with 10ug/mL cytochalasin D, 10ug/mL LPC, 100ug/mL cycloheximide, 1uM nocodazole, energy mix (7.5 mM creatine phosphate, 1 mM ATP pH 7.7, 1 mM $MgCl_2$) at 1:100 dilution. To monitor the quality of the nucleus and label the cytoplasm in the filtration experiment, ~1ug/mL Hoechst dye, ~0.5uM NLS-GFP, and 1:1000 dilution of MitoTracker Red (M7512 ThermoFisher) were added to the extract. The extract was stored on ice for further use.
To isolate the nuclei from the cytoplasm we used 3D filter holders. The design files are available on our GitHub page (https://github.com/wuhrlab/3DFilterHolderDesigns). We used the Hubs platform (https://www.hubs.com/) to print the holders using either Standard Resin (SLA) or Dental resin (SLA) materials at 20% infill rate, 50µm layer height.
The undiluted embryonic cell lysate was filtered through polycarbonate membranes with uniform pore sizes (5µm) using a table-top centrifuge at 2,000g for 2.5min at 4°C. Nuclei remained in the supernatant while nuclear-depleted cytoplasm flowed through the membrane. To further remove cytoplasmic impurities, the supernatant was diluted 2-fold with XB buffer, and the filtration spin was repeated at 2,000g for 2.5min at 4°C. The two flow-through fractions were collected and combined. Lysates were incubated with Hoechst, NLS-GFP, and MitoTracker for checking via imaging. At each stage, the nuclear and cytoplasmic fractions were digested into tryptic peptides, labeled with isobaric tags, and subjected to accurate multiplexed proteomics analysis, as described.
Canonical nuclear proteins are defined as proteins that are classified as nuclear in the frog oocyte[4] and various other cell types from published databases such as hyper LOPIT[28], Cell Atlas[29], Protein Atlas[30], and Uniprot[31]. To avoid the fraction of proteins are promiscuously assigned to several subcellular localizations, the list excluded proteins whose assigned localizations also include the following compartments: mitochondria, Golgi, endoplasmic reticulum,cytosol, cytoskeleton, plasma membrane.
To correct for cytoplasmic proteins retained on the filter, we subtracted the signal of the 2-cell stage supernatant experiment (negligible nuclear amounts) from all other supernatant experiments. From

the MS analysis of nuclear filtration assays, the relative protein signal in each fraction at each collected point was obtained.Using the immunofluorescence (IF) data, we converted the time post fertilization to the nuclear-to-cell volume ratio (NCV-ratio) using the "slmenginefunction" from MathWorks File Exchange, created by John D'Erric. For each protein, the nuclear fraction was fitted by a sigmoidal function of NCV-ratio assuming the final nuclear fraction value reached the value in the oocyte[4]. The fit parameter (bound by 0 and 1) was defined as the NCV-ratio value when 50% protein amount enters the nucleus. Using the same spline fit of the IF data, this ratio was converted to $T_{embryo1/2}$, which is defined as the time post fertilization at 16°C when 50% protein enters the embryonic nuclei.

Nuclear isolation with a commercial kit (Abcam ab113474)

Embryo extract at cleavage 12 (the ZGA) was prepared as described above. The undiluted embryonic extract was projected to the Abcam Nuclear Extraction Kit (ab113474) and followed the manufacturer's suggested protocol. The resulting nuclear and cytoplasmic fractions were digested into tryptic peptides, labeled with isobaric tags, subjected to the MS, and nuclear fraction analysis as described above.

MS sample preparation and analysis

Samples were prepared mostly as previously described[32]. Lysates were collected in 100 mM HEPES pH 7.2. To reduce disulfides, Dithiothreitol (DTT) (500 mM in water) was added to a final concentration of 5 mM (20 min, 60°C). Samples were cooled to RT, and cysteines were alkylated by the addition of N-ethyl maleimide (NEM, 1 M in acetonitrile) to a final concentration of 20 mM followed by incubation for 20 min at RT. 10 mM DTT (500 mM stock, water) was added at RT for 10 min to quench any remaining NEM. A methanol-chloroform precipitation was performed for protein clean-up, and the collected protein pellets were allowed to air dry. Samples were taken up in 6M guanidine chloride in 200 mM EPPS pH 8.5. Subsequently, the samples were diluted to 2 M guanidine chloride in 200 mM EPPS pH 8.5 for overnight digestion with 20ng/µL Lys-C (Wako) at RT. The samples were further diluted to 0.5 mM guanidine chloride in 200 mM EPPS pH 8.5 and then digested with 20 ng/µL Lys-C and 10 ng/µL trypsin (Promega) at 37°C overnight.

The digested samples were dried using a vacuum evaporator at RT and taken up in 200 mM EPPS pH 8.0. Then total material from each condition was labeled with tandem mass tags (as indicated by the experiment: TMT-6plex, TMT-11plex, TMTpro-16plex - Thermo Fisher Scientific). TMT/TMTpro samples were labeled for 2 hours at RT. Labeled samples were quenched with 0.5% hydroxylamine solution. Samples from all conditions were combined into one tube, acidified to pH < 2 with phosphoric acid (HPLC grade, Sigma) and cleared by ultracentrifugation at 100,000 rcf at 4°C for 1 hour in polycarbonate tubes (Beckman Coulter, 343775) in a TLA-100 rotor. Supernatants were dried using a vacuum evaporator at RT. For a low complexity sample, dry samples were taken up in HPLC-grade water and stage-tipped for desalting[33] and resuspended in 1% formic acid (FA) to 1 µg/µL for mass spectrometry analysis. For high complexity samples, the supernatant was sonicated for 10 minutes and then fractionated by medium pH reverse-phase HPLC (Zorbax 300Extend C18, 4.6 x 250 mm column, Agilent) with 10 mM ammonium bicarbonate, pH 8.0, using 5% acetonitrile for 17 minutes followed by an acetonitrile gradient from 5% to 30%. Fractions were collected starting at minute 17 with a flow rate of 0.5 mL/min into a 96 well-plate every 38 seconds. These fractions were pooled into 24 fractions by alternating the wells in the plate[34]. Each fraction was dried and resuspended in 100 µL of HPLC water. Fractions were acidified to pH <2 with HPLC-grade trifluoroacetic acid, and stage-tipping was performed to desalt the samples. For LC-MS analysis,

samples were resuspended to 1 µg/µL in 1% FA and HPLC-grade water, and ~1 µg of peptides were analyzed per 1 hour run time.
Approximately 1-3ug of the sample was analyzed by LC-MS. LC-MS experiments were performed with an nLC-1200 HPLC (Thermo Fisher Scientific) coupled to an Orbitrap Fusion Lumos (Thermo Fisher Scientific). For each run, peptides were separated on an Aurora Series emitter column (25cm x 75µm ID, 1.6µm C18) (ionopticks, Australia), held at 60°C during separation by an in-house built column oven. Separation was achieved by applying a 12% to 35% acetonitrile gradient in 0.125% formic acid and 2% DMSO over 90 min for fractionated samples and 180 min for unfractionated samples at 350 nL/min at 60°C. Electrospray ionization was enabled by applying a voltage of 2.6 kV through a MicroTee at the inlet of the microcapillary column. As indicated in each proteomics experiment, we used the Orbitrap Fusion Lumos with a TMT-MS3[35], TMTc+[36], TMTpro-MS3[37], or TMTproC[38] as previously described. The mass spectrometry proteomics data have been deposited to the ProteomeXchange Consortium via the PRIDE partner repository with the dataset identifier PXD02806[39]. To access, please use username:, and password: Iw5b6KTw.
Mass spectrometry data analysis was performed essentially as previously described[36]. The mass spectrometry data in the Thermo RAW format was analyzed using the Gygi Lab software platform (GFY Core Version 3.8) licensed through Harvard University. Peptides that matched multiple proteins were assigned to the proteins with the greatest number of unique peptides. TMT-MS3 [35], TMTc+ [36], TMTpro-MS3 [37], or TMTproC [38] data were analyzed as previously described.

RNA gel analysis

The RNA gel image was extracted from Newport and Kirschner's 1982 publication[17] and analyzed using ImageJ. The relative intensity of each RNA band was measured 5 times and the mean value was reported. For snRNA, there were 5 visibly distinct bands that were each measured individually.

Importin affinity assay

For importin and Ran constructs, we received plasmids gift for the following constructs: GST-importin α (*Xenopus*) and GST-importin β (*Xenopus*) from Sabina Petry, His-Tev- RanQ69L and ZZ-importin β (*Homo sapiens*) from Dirk Görlich and Thomas Güttler. Proteins were expressed and purified mostly as previously described[40,41]. Briefly, all constructs were transformed into Rosseta2 E. coli cells (Fisher: 71-403-4) for protein expression and were grown in TB media (Sigma: T0918) prepared according to the supplier's instructions. Cells were grown at 37˚C, shaking at 200 RPM. When an OD600 of 0.8 was reached, cells were induced with 0.15 mM isopropyl-β–D-1- thiogalactopyranoside (IPTG) and grown overnight at 20˚C 200 RPM to reach an OD600 post-induction of 25.6. The culture was harvested the next morning by centrifugation at 4°C at 5,000g with Beckman J2-MI centrifuge with JA-10 rotor for 20 min. Cells were lysed on ice using 0.25mg/mL lysozyme in lysis buffer (50mM K-phosphate pH 7.0, 500mM NaCl, 5mM $Mg(OAc)_2$, 1mM Ethylenediaminetetraacetic acid (EDTA), 2mM DTT, 40U/mL Benzonase Nuclease (Novagen 70746-4) and 2mM PMSF for 10 min. After lysozyme digestion, the lysate was pipetted up and down until a homogenous mixture was reached. Cells were then further homogenized using an EmulsiFlex (Avestin) in lysis buffer. The lysate was clarified, and the supernatant was collected. Most of the constructs continued to further purification steps, except for ZZ-tag-Importin-β, which was aliquoted in 10-20µL aliquots, flash-frozen using liquid $N_2$, and stored at -80C for the importin interaction experiment.
For GST-importin α, and GST-importin β constructs, the lysates were bound to Pierce Glutathione Agarose resin (ThermoScientific 16101), washed, and eluted in lysis buffer containing 10mM Glutathione. The eluents were collected for further purification.

For His-Tev-RanQ69L construct, the lysate was bound to NiNTA agarose beads (Qiagen 1018236), washed in lysis buffer + 10uM Guanosine-5'-Triphosphate Disodium Salt (GTP) (ThermoScientific 56001-37-7), and eluted in lysis buffer containing 200 mM Imidazole + 10uM GTP. The collected eluent of His-Tev-RanQ69L was then subjected to an overnight His-TEV protease (Invitrogen 10127-017) cleavage at 10U per 100ug target proteins and dialysis sequence to exchange to final buffer solution of CSF-XB (100mM KCl, 20mM HEPES, 2mM $MgCl_2$, 0.1mM $CaCl_2$, 4mM Ethylene glycol-bis(2-aminoethylether)-N,N,N′,N′-tetra-acetic acid (EGTA), pH7.8) + 30uM GTP. The dialyzed solution was bound to the 2$^{nd}$ NiNTA column to remove His-TEV protease, and the Ran construct was eluted in the final buffer for further purification.

All protein constructs, after elution, were further purified using gel filtration (Superdex 200 HiLoad 16/600, GE Healthcare – 28-9893-35) in CSF-XB buffer and 250mM sucrose. The purity of the proteins was confirmed by Coomassie-stained SDS-PAGE gels. Protein concentration was determined using an A280 Nanodrop (Thermo Scientific Nanodrop lite) with the corresponding extinction coefficient (calculated based on protein sequence and using ProtParam calculator https://web.expasy.org/protparam/). 10-20μL protein aliquots were flash-frozen using liquid $N_2$ and stored at –80C.

In the importin testing system, both crude interphase extract and clarified extract were used. Clarified extracts were prepared as described[42]. Briefly, crude mitotic extracts were spun for a second time at 50,000 rpm in a Beckman TLS-100A rotor for 2hr at 4°C. The clear middle layer was extracted using a 22-gauge needle. Fresh interphase egg extract was made as described earlier.

In experiments using GST-importin-β, Pierce Glutathione Magnetic Agarose beads (Thermo Scientific 78602) were washed twice in XB buffer and then incubated in 1 hr with purified GST-importin-β at 2μg/μL proteins per μL beads. Flow-through importin β was removed post incubation to avoid free importin β competing with the immobilized β in extract. GST-importin-α was added at the ratio 1:1 importin α: importin β molar concentration and incubated for 30 min. RanQ69L was titrated into the solution at estimated importin β:RanQ69L ratios of 1:0 to 1:100. Finally, extract was added to the mixture so that the final importin β concentration reached 10μM and incubated for 1 hour at 16°C. After the incubation, beads were washed two times with XB buffer and eluted with sample buffer (Invitrogen NP0007) for a quick check with Coomassie gels (Invitrogen NW00100BOX) and eventually MS analysis.

In experiments using ZZ-tag-importin-β construct (expressed as described earlier), IgG Sepharose 6 Fast Flow (Cytiva 17-0969-01) were washed twice in XB buffer and then incubated with the cell lysate of the overexpressed construct for 1 hr. The supernatant was then removed, and beads were washed once with XB buffer. RanQ69L was titrated into the solution at an estimated RanQ69L ratio of 1:0 to 1:100 and incubated for 30 min before removing the flow-through. GST-importin- β was added at the ratio 1:1 molar concentration. Finally, the fresh extract was added, and the pull-down collection was performed as described above.

The relative protein signals from the MS analysis were normalized by the added importin signal and IgG signals (in the case of the ZZ-tag-importin-β experiment). The protein fraction was defined as signal of importin-bound protein in a condition with RanQ69L divided by the sum signal of protein in the conditions with and without RanQ69L, reflecting the change in the RanQ69L amount in each condition. The protein fractions were then normalized using the values of known background proteins (such as glycolytic enzymes and mitochondrial proteins) that do not interact with importin β. For each protein, the fractions were fitted through a linear function of the normalized added RanQ69L amount detected in the pull-downs. The y-intercept was fixed at 0.5. The extracted slope was used as a proxy for the protein's affinity to importin β. The experiments were repeated three times, and the measurements were projected onto a single dimension that maximized the agreement in variation between the $T_{embryo1/2}$ and the importin affinity proxy using canonical correlation analysis[18]. The projection is cross validated as following: The dataset was split into 10 consecutive folds with shuffling. At each fold, 90% of the data was used for training the canonical correlation axis while 10% was kept behind for validation. The validated values of the 10% portion from each round were collected from each round and made up the final vector of cross-validated projected values on the canonical axis. The projected values defined the final proxy for importin affinities and were used for downstream analysis.

DNA affinity assay

A pQE-80L empty plasmid (from SnapGene) was cut using EcoRI and BamHI (New England Biolabs) and purified using QIAquick Gel Extraction Kit (Qiagen). The collected DNA fragments were end-filled using Klenow (New England Biolabs) and biotin-dATP (Invitrogen). DNA was coupled to streptavadin Dyna beads (65305; Invitrogen) following the protocol outlined previously[43]. DNA bound to beads was collected at the estimated 1µg/µL per beads volume and saved for future experiments.

Fresh interphase egg extract was made as described earlier. DNA beads were incubated in fresh extract for 2 hours at concentrations ranging from 60ng/µL to 160 ng/µL. To arrest extract in the interphase, cycloheximide was added along with Hoechst and NLS-GFP to monitor the formation of a nucleus with epifluorescence microscopy. Post incubation, pull-downs were collected and eluted with 6M Guanidine chloride pH 7.2 and subjected to MS analysis.

Absolute Protein Concentration Estimates

For absolute protein abundance estimates (Supplementary Table 4), we reanalyzed previously collected mass spectrometry data with a protein reference database based on the 9.2 version of the *Xenopus laevis* genome downloaded from Xenbase (http://ftp.xenbase.org/pub/Genomics/JGI/Xenla9.2/sequences/XENLA_9.2_Xenbase.pep.fa)[19,44-46]. The previously published estimates were analyzed with an mRNA reference database. To deduce the power-law relationship between MS-signal and protein concentration, we generated using a regression linking the average log ion signal per peptide to pre-existing estimates of the protein concentration[19,44]. We measured the ion flux integrated over time by peptide, determined the total log ion signal for the protein, and divided by the total number of peptides in the protein to calculate the average log ion signal per peptide. We related the normalized log ion signal to the pre-existing protein concentration estimates using a robust regression since such data is often error-prone and skewed by outliers. We binned normalized log ion signals into bins of size 1/3 (in log ion signal space), calculate the median protein concentration across all proteins with normalized log ion signals in the range of the bin, and then fit a robust regression with a trimmed mean M-estimator with Ramsay's Ea of 1.65. We use that fit to estimate the protein concentration for all proteins detected in the *Xenopus* egg through mass spectrometry.

Modelling of nuclear import in embryos and encapsulated droplets

We develop a model for nuclear import in early embryos based on the differential affinities of proteins to importin and the experimentally observed increase in total nuclear volume.

In this simple description, an embryo with a constant total volume ($\mathrm{V_{embryo}}$) contains protein $i$ , where $i = 1, \dots, n$ for $n$ total proteins, with $P_i$ is protein abundance of $i$ with an embryonic concentration $[\mathrm{P_i}] = \frac{\mathrm{P_i}}{\mathrm{V_{embryo}}}$. Protein $i$ is imported from the cytoplasm to the nucleus, where $[\mathrm{P_{cyto,i}}] = \frac{\mathrm{P_{cyto,i}}}{\mathrm{V_{embryo}}}$ and $[\mathrm{P_{nuc,i}}] =$

$\frac{P_{nuc,i}}{V_{embryo}}$ represent embryonic concentration of protein $i$ located in the cytoplasm and nucleus respectively. Here, $P_{cyto,i}$ is the abundance of $i$ in cytoplasm and $P_{nuc,i}$ is in the nucleus. Protein $i$ undergoes the following transformation:

$$P_{cyto,i} \rightarrow P_{nuc,i} \quad (1)$$

Note that $[P_{cyto,i}] + [P_{nuc,i}] = [P_i]$ to mass balance equation #1. If we denote the net flux of proteins from the cytoplasm into the embryonic nuclei as $F_{total}$, for a particular protein $i$ we can write the change in its abundance of those resided in the nucleus of the embryo as:

$$\frac{dP_{nuc,i}}{dt} = \theta_i F_{total} \quad (2)$$

Where $\theta_i$ is the fraction of the total nuclear import flux contributed by protein $i$. The total nuclear import flux $F_{total}$ is the rate at which the total embryonic nuclear volume increases throughout development multiplied by the total protein concentration in the embryos.

$$F_{total} = \frac{dV_{nuc}}{dt}\sum_{i=1}^{n}[P_i] \quad (3)$$

Based on experimental result, the total protein concentration $\sum_{i=1}^{n}[P_i]$ stays roughly constant throughout early development and is approximately ~2 mM (Supplementary Table 1)[19]. The rate of nuclear volume expansion, $\frac{dV_{nuc}}{dt}$ is written as the time derivative of the total nuclear volume as derived from our immunofluorescence data (Extended Data Fig. 5a). We assume that 70% of the cytoplasmic volume is excluded by yolk and lipids[4,47].

To derive $\theta_i$, the fraction of nuclear influx contributed by protein $i$, we assume that substrate binding to importin is in equilibrium, as in the Langmuir adsorption isotherm model[48].

$$P_{cyto,i} + I \rightleftarrows^{K_{Di}} IP_{cyto,i} \quad (4)$$

Where $[I]$ is the importin concentration in the embryo, which has a fixed total concentration $[I_o]$ throughout the considered developmental period[19], and $K_{Di}$ is the equilibrium dissociation constant for protein $i$ in this reaction. At equilibrium, $K_{Di}$ is:

$$K_{Di} = \frac{[P_{cyto,i}][I]}{[IP_{cyto,i}]} \quad (5)$$

Using the mass balance equation

$$[I] = [I_0] - \sum_{i=1}^{n}[IP_{cyot,i}] \quad (6)$$

we can combine equations 5 and 6 to find the fraction of total importin that is bound by protein $i$:

$$\theta_i = \frac{[IP_{cyto,i}]}{[I_o]} = \frac{\frac{[P_{cyto,i}]}{K_{Di}}}{1 + \sum_{j=1}^{n}\frac{[P_{cyto,j}]}{K_{D,j}}} \quad (7)$$

Combining equations 2, 3, and 7, we arrive at a system of $n$ differential equations, where $n$ is the total number of proteins:

$$\frac{dP_{nuc,i}}{dt} = \left(\frac{\frac{[P_{cyto,i}]}{K_{Di}}}{1 + \sum_{j=1}^{n}\frac{[P_{cyto,j}]}{K_{Dj}}}\right)\frac{dV_{nuc}}{dt}\sum_{k=1}^{n}[P_k] \quad (8)$$

This system of $n$ differential equations and mass balance equations are solved using the ode45 function in MATLAB.

We created a synthetic system of nuclear proteins over a distribution of importin affinities in a background of importin-inert cytoplasmic proteins to mimic an actual embryo. We estimate that nuclear proteins constitute ~10% of the embryonic proteome based on the quantified NCV-ratio in the oocyte (Fig. 1b). Nuclear proteins' affinities to importin are sampled from a log-normal distribution, i.e. the natural log of $\mathrm{K_{Di}}$ are sampled from $\ln\,(\mathrm{K_{Di}}) \sim \mathrm{N}(\mathrm{mean} = -18, \mathrm{standard\ deviation} = 2)$ which corresponding to a median affinity ($\mathrm{K_D}$) of ~30nM - a benchmark binding affinity measured for protein-protein interactions[49,50]. Cytoplasmic proteins account for 90% of the proteome and have no affinity to importin ($\mathrm{K_{Di}} = \infty$). We estimate the protein concentration for each species from the total protein abundance estimation in the egg[19]. A median protein concentration is ~44nM and the total protein concentration is ~2mM in the egg (Supplementary Table 4)[19].

We simulated embryonic nuclear import with 4,600 nuclear proteins, for each at a concentration of 44nM representing 10% of the proteome, cytoplasmic proteins at 1.8mM concentration and 1.5µM of importin (constant over time). For an illustration of differential nuclear entry due to importin affinities, two representative proteins (one with high importin affinity (10nM) and one with low importin affinity (1µM)) show the expected differential nuclear entry times into the increasing embryo's nuclear volume (Fig. 3e). For an illustration of sequential titration of nuclear concentration due to the continuous nuclear import over development, we simulate three representative proteins: a high affinity protein (1nM), an intermediate affinity (30nM), and a low affinity (1µM).

Similarly, we test the model in a single cell cycle after the reformation of nuclear envelop. Compared to the embryo simulations we adapted a modification on nuclear flux: The nuclear flux, $\frac{\mathrm{dV_{nuc}}}{\mathrm{dt}}$ is derived from the change of nuclear volume in cell-free droplets (Extended Data Fig. 5c). We simulate a similar system as in early development with the adjusted nuclear flux and illustrated that nuclear concentration of titrating proteins from 1nM, 30nM, to 1µM, sequentially reaches the maximum as the nucleus grows in cell droplets (Fig. 4e).

Assaying nuclear import in oil encapsulated artificial cells

The Gateway entry plasmids of desired proteins were retrieved from *Xenopus laevis* ORFeome [51]. The destination vector carrying an EGFP sequence – TEV site – S-tag (pCSF107mT-GATEWAY-3'-LAP tag) was chosen and bought from Addgene. For the Gateway LR cloning reaction, the entry plasmid, the destination plasmid, and the Gateway LR clonase II enzyme mix (Invitrogen 11791) were combined at the ratios recommended in the manufacture protocol. After the reaction, the expression cloned vector was purified, then linearized using restriction enzymes, which were chosen so that the region of protein of interest was protected. The linearized plasmids were *in-vitro* transcribed using the mMESSAGE mMACHINE SP6 kit (Invitrogen AM1340) supplemented with a 7-methyl guanosine cap protected on the 5' end terminal, and a poly(A) tail (NEB M0276). Finally, RNA products were purified using Trizol LS reagent (Invitrogen 10296010), then resuspended in nuclease-free water at ~1ug/µL in the final RNA concentration.

We microinjected the RNA products into the oocytes using a PM2000B 4-channel Pressure Injector (MicroData Instrument). ~100 oocytes per protein construct were injected twice to be equally distributed around the animal cap at the total volume of ~50 nL. Injected embryos were allowed to recover in 2.5% Ficoll OCM and visually inspected before use in all experiments. After 1-hour resting in Ficoll, oocytes were transferred to OCM for overnight expression. Only healthy oocytes were used the next day to make the extract. Oocyte extract was collected as previously described. Finally, the undiluted extract was supplemented with 50mM sucrose, 20ug/mL LPC protease inhibitors, and 20ug/mL cytochalasin D. The presence of the desired protein was validated via epifluorescence imaging of anti-S-tag beads pulled down from the extract (SinoBiological MB101290-T38). After confirming the expression, the extract was flash-frozen and stored in ~1µL aliquots at -80C for further use.

CSF-arrested *Xenopus laevis* egg extract was driven into interphase by supplementing $Ca^{2+}$ [4mM] and demembranated sperm nuclei[52], which served as chromatin sources for nuclear assembly, to a final concentration of 1E6 nuclei/µL. To facilitate imaging of growing nuclei, mCherry GST-NLS 11.5 mg/mL was added to the master mix at a 1:100 dilution. Candidate GFP-protein conjugates were also added to the extract mix at 1:25 dilution prior to nuclei encapsulation in droplets, with concentrations ranging from 0.5mg/mL – 3.0 mg/mL depending on the protein being used.

Polydimethylsiloxane (PDMS) T-junction microfluidic devices affixed to #1.5 coverslips were used to generate monodisperse emulsions of ~50 µm diameter extract droplets in a continuous oil phase as previously described[53]. To facilitate the use of small extract volumes, 2mL Eppendorf tubes containing 25 µL of extract were placed on ice and pressurized using microfluidic pressure pumps (Flow EZ 1000, Fluigent). Flowrates of both the extract and oil phases were controlled by modulating the applied pressures. PDMS devices were kept on ice during filling. Once a device was filled, its inlet and outlet channels were sealed with silicon tubing plugs and the device was then placed on the microscope stage for imaging.

All imaging was performed in a temperature-controlled room at 18°C. Image acquisition was conducted using an IX-81 confocal microscope (Olympus USA) equipped with a 40x 0.6NA objective and a CSU-W1 confocal scanning unit. Images were captured with an ORCA-Flash4.0 sCMOS camera (Hamamatsu Photonics) and system automation was controlled via cellSens software (Olympus USA). Time-lapse image series of protein import into encapsulated nuclei were generated by acquiring z-stacks (step-size = 2 µm, 12 slices per time point) in both the red and green channels at 2 min intervals. Acquisition began 5-6 minutes after filled devices were removed from ice (we called this t = 0) and typically lasted for 1-2hr.

The collected image series were analyzed using Fiji software (NIH). Once collected, hyperstacks were parsed by channel and a modified version of the Autofocus hyperstack macro (Richard Mort) was used to extract a single, best-focused image of the nucleus for each time point and generate a time-lapse series of nuclear import. Before proceeding with analysis, automated best-focused selections were then confirmed manually. Images were typically segmented using the mCherry signal as a reference to generate ROIs outlining the nucleus. In the rare case in which the GFP-protein entered the nucleus first, the GFP channel was used as a reference to generate ROIs. These ROIs were then copied to corresponding images in the opposing channel and the mean and integrated fluorescence intensities of the nuclear ROIs were measured for each channel, red and green, using the Analyze Particles function in Fiji. To measure cytoplasmic fluorescence intensities, nuclear ROIs were dilated, and the same measurements repeated. Cytoplasmic intensity was then calculated by subtracting the integrated intensity of the original nuclear ROI from the dilated ROI integrated intensity. The relative nuclear concentration was calculated by dividing the mean nuclear intensity by the sum of the mean nuclear and cytoplasmic intensities. Since the extract was well-mixed and uniform before the formation of the nucleus, the RNC at the initial time point was 0.5. The RNC data were fitted with a sigmoid function to extract the time $T_{droplet\ 1/2}$, at which the relative intensity reaches half of its max value for each protein and the corresponding mCherry-NLS. To overcome extract variability, the import time difference ($\Delta T_{droplet\ 1/2}$) between mCherry-NLS and the GFP-conjugated protein of interest was used to compare the nuclear import rates between proteins of interest.
